# Evidence of learning walks related to scorpion home burrow navigation

**DOI:** 10.1101/2021.12.28.474378

**Authors:** Douglas D. Gaffin, Maria G. Muñoz, Mariëlle Hoefnagels

**Affiliations:** Department of Biology, University of Oklahoma, Norman, OK 73019 USA; Department of Microbiology and Plant Biology, University of Oklahoma, Norman, OK 73019 USA

**Keywords:** pectines, peg sensilla, familiarity, sensory

## Abstract

The *Navigation by Chemotextural Familiarity Hypothesis* (NCFH) suggests that scorpions use their midventral pectines to gather chemical and textural information near their burrows and use this information as they subsequently return home. For NCFH to be viable, animals must somehow acquire home-directed “tastes” of the substrate, such as through path integration (PI) and/or learning walks. We conducted laboratory behavioral trials using desert grassland scorpions (*Paruroctonus utahensis*). Animals reliably formed burrows in small mounds of sand we provided in the middle of circular, sand-lined behavioral arenas. We processed overnight infrared video recordings with a MATLAB script that tracked animal movements at 1-2 s intervals. In all, we analyzed the movements of 23 animals, representing nearly 1500 hours of video recording. We found that once animals established their home burrows, they immediately made one to several short, looping excursions away from and back to their burrows before walking greater distances. We also observed similar excursions when animals made burrows in level sand in the middle of the arena (i.e., no mound provided). These putative learning walks, together with recently reported PI in scorpions, may provide the crucial home-directed information requisite for NCFH.

## INTRODUCTION

Sand scorpions live in burrows that they dig, and from which they emerge at night to hunt (Polis and Farley, 1979; Polis, 1980; Polis and Farley, 1979; Polis et al., 1985). Questions exist about how they return home. We think scorpions might use a simple view-based navigational process, similar to that proposed for ants and bees, termed *Navigation by Scene Familiarity* (Baddeley et al., 2012; Philippides et al., 2011). However, instead of vision, scorpions may be guided by tastes and touches acquired via their mid-ventral pectines (Cloudsley-Thompson, 1955; Wolf, 2017). We have termed this process *Navigation by Chemo-textural Familiarity* (Gaffin and Brayfield, 2017). Put simply, to get home, the scorpion uses its pectines to detect and move toward tastes and textures it has learned during previous home-bound forays.

For this hypothesis to be viable, two crucial ingredients must be present. First, there must be adequate sensor complexity to match the environment. Second, there must be a way to generate the initial home-bound training paths (Baddeley et al., 2012; Gaffin et al., 2015; Wehner et al., 1996).

Regarding sensor complexity, each pecten has a series of teeth that support thousands of minute peg sensilla (aka “pegs”) on their ground facing surfaces (Ivanov and Balashov, 1979; Foelix and Müller-Vorholt, 1983). Each peg contains a population of chemosensory taste cells (approx. 10) and at least one mechanosensory neuron that responds when the peg bends (Ivanov and Balashov, 1979; Foelix and Müller-Vorholt, 1983; Gaffin and Brownell, 1997b; Melville, 2000). In all, hundreds of thousands of sensory afferents project from the pectines to the scorpion’s central nervous system (Wolf, 2008; Brownell, 2001; Drozd et al., 2020). Based on this complexity, a proof-of-concept model showed that an agent using a downward facing sensor could navigate various proxies of a simulated environment (Musaelian and Gaffin, 2020).

Sensory complexity is therefore adequate; what about the generation of home-bound training paths? The glances and tastes a scorpion experiences while leaving its nest or burrow depart 180 degrees from those that lead home (Fig 1). How does the animal know its way home after venturing out for the first time? Innate behaviors such as path integration and learning walks may provide the answer. In path integration (PI), the distance and direction of each outbound leg is integrated to compute an approximate homebound vector (Wehner, 1992; Papi, 1992). PI is well documented for many animals, but the studies of desert ants are the most extensive (Wehner, 1992; Wehner and Srinivasan, 1981; Wehner et al., 1996; 2004; 2006; Wolf, 2011; Wittlinger et al., 2006; 2007; Wittlinger and Wolf, 2013; Heinze et al., 2018; Srinivasan, 2015; Wittlinger et al., 2006; Wolf, 2011). PI has also been described for some groups of spiders (Ortega-Escobar, 2002; 2006; Ortega-Escobar and Ruiz, 2017; 2014; Görner and Claas, 1985; Moller and Görner, 1994; Seyfarth and Barth, 1972; Seyfarth et al., 1982), and a recent study showed evidence of path integration in the lesser Asian scorpion, *Mesobuthus eupeus* (Prévost and Stemme, 2020).

**Fig. 1.**
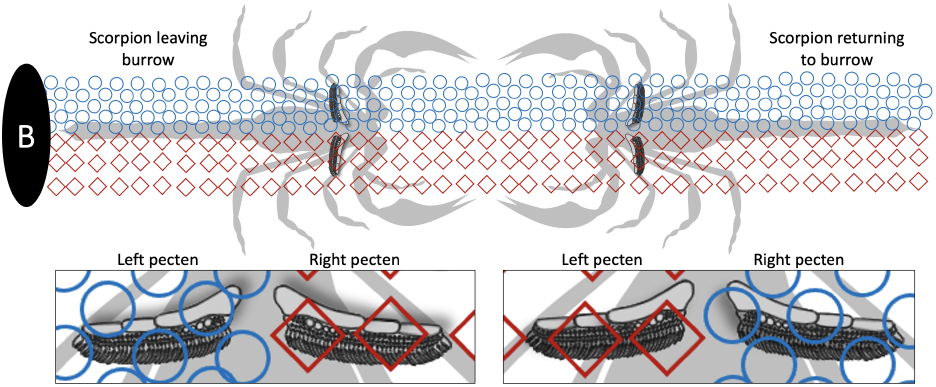
Conflicting information between outbound vs inbound paths. The chemicals and textures the pectines experience on the journey leading away from the burrow depart 180 degrees from what they experience on the return trip to the burrow.

In addition, learning walks are innate behavioral patterns thought to further help the animal gain goal-directed stimuli (Wehner et al., 2004; Fleischmann et al., 2016; Zeil et al., 2014; 1996). As with PI, learning walks are well described for navigating ants (Zeil and Fleischmann, 2019) but have never been documented for scorpions or any other arachnid (although the zigzag outbound paths of the wandering Namib spider *Leucorchestris arenicola* are strongly suggestive (Narendra et al., 2013; Nørgaard et al., 2012; Gaffin and Curry, 2020)).

In this study, we made long-term video recordings of sand scorpions as they produced burrows in the middle of laboratory arenas. We show that the animals make consistent, repeated looping paths immediately after their first burrow digging behavior and that these paths have similar characteristics to learning walks in ants.

## MATERIALS AND METHODS

### Animals, collection details, and maintenance

Desert grassland scorpions (*Paruroctonous utahensis*) were collected from the Walking Sands dune area about 6 km SE of the UNM Sevilleta Field Station. We used UV lights to find animals on three nights during periods of new moon in August, September, and October 2020. Only animals judged to be adults were collected. Supplemental Fig 1 shows the collection locations and the mixture of males and females from the three collection nights. Males dominated the August and September collections, whereas females predominated in October. The animals were transported and housed individually at the Station in small rectangular food storage containers with air holes drilled in the lids and ∼50 ml of sand collected from the animals’ habitat as a substrate. The animals were exposed to a 14:10 h light:dark cycle (on at 06:00, off at 20:00) using indirect light from two white 60W equivalent LED bulbs housed in work lights (Bayco clamp light, 21.6 cm) placed ∼50 cm from the animals and plugged into a timer switch. The room temperature was maintained at about 22°C. After 45 days, we moved all animals to a room in the laboratory building on the UNM Sevilleta campus where the animals were exposed to natural light that streamed through the large NE facing picture windows and the temperature was kept at 20-21°C and the RH between 16-20%. A voucher specimen was given to the Sam Noble Oklahoma Museum of Natural History at the University of Oklahoma in Norman, Oklahoma.

### Encouraging burrow formation

We ran several pilot studies to determine which conditions were most conducive to the scorpions digging and occupying burrows. These included tests of various substrates, mound sizes and locations, sand moisture content, and the timing of burrow occupation relative to daylight. All of these tests are chronicled in the supplemental materials section (Supplemental Figs 2-7).

### Behavioral apparatus and video recording

We built four identical behavioral set-ups (Fig 2) in the UNM Sevilleta field station lab building (inspired by Vinnedge and Gaffin (2015)). Each arena consisted of an aluminum water heater drain pan (Camco, product no. 20860; 76.6 cm base diam, 7.6 cm height) sitting atop a turntable (formed from a 70 cm diam x 1.9 cm thick plywood round attached to a 30.5 cm diam Richelieu swivel plate with 454 kg capacity) to allow 360° rotation. A rubber mat (Ottomanson multi-purpose 61 × 61 cm exercise tile mat) was placed beneath each arena to dampen room vibrations. About 1250 ml of screened native sand was spread in a thin layer across the bottom of each arena. We then added ∼250 ml of native sand through a small funnel to form a mound in the middle of the arena which we then misted from above with 20 squirts of water (∼15 ml). Light blocking curtains were secured to hula hoops (Ice Hoop, Kess Co; 86 cm diam) with large binder clips and draped around each arena. Two work lights (Bayco clamp light, 21.6 cm) equipped with broad spectrum bulbs (Duracell Ultra 75W equivalent daylight) were positioned 110 cm above each arena. The lights were controlled by a timer set to a 14:10 h light:dark cycle (on at 0530, off at 1930). Infrared cameras (ELP 1 megapixel Day Night Vision) were mounted 110 cm above the centers of each arena and connected via USB to two laptop computers (two cameras per laptop; Apple MacBooks). A MATLAB script was written to toggle between the cameras and acquire 200×200 pixel frames at a user defined interval. The frames were stored in a MATLAB structure array for subsequent analysis.

**Fig. 2.**
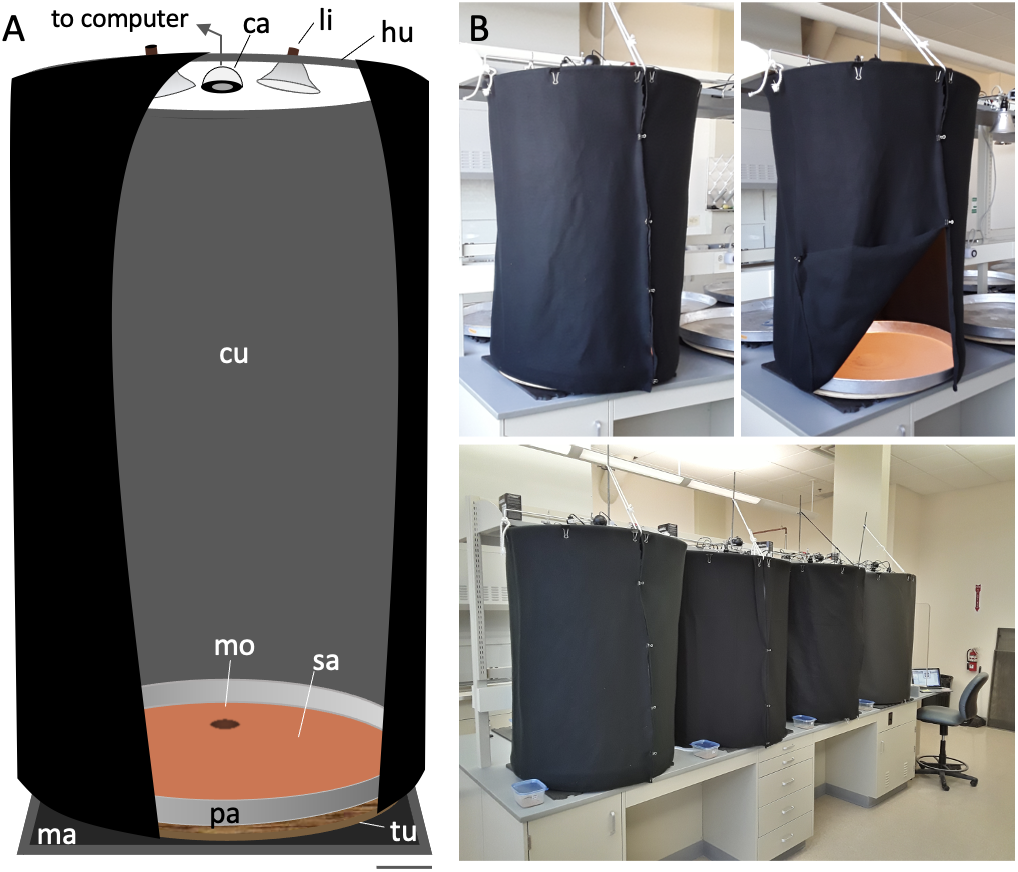
Behavioral set-up for long-term recordings. (A) Each arena was composed of an aluminum heater drain pan (pa) atop a turntable (tu) and a rubber mat (ma). Sand (sa) was added to the pan and a mound (mo) was formed in a pre-defined location. A curtain (cu) was cut from black, light-blocking material and suspended from a hula hoop (hu) attached to supporting frame. Two timer-controlled lights (li) and an IR camera (ca) connected to a laptop computer were also attached to the frame. (B) Photos of an arena with curtain closed (left) and open (right) and the four arenas (below) arrayed along the lab counter with the laptop computer at the end.

To aid in video tracking, we used double-sided tape to affix a small crystal (5 mm round cab crystal; Acrylic Gems) on the dorsal mesosoma of each animal before releasing them into the behavioral arenas (Fig 3). To secure the crystal, we first placed an animal in a rectangular plastic container (30 × 17 cm). We then placed a square plastic sheet (8.5 × 8.5 cm) that had a 6 mm hole cut close to one of its corners over the animal such that the hole was over the animal’s mesosoma with the remainder of the sheet covering the rest of the animal’s body. This system calmed and secured the animal and allowed the crystal to be easily applied through the hole to the animal’s back without the danger of being stung. The crystals reflected IR from all angles and from all animal positions within the arenas, so the plotting accuracy in MATLAB was greater than 99%. Smaller 3 mm crystals proved less effective, given the camera’s resolution and distance from the arena floor.

**Fig. 3.**
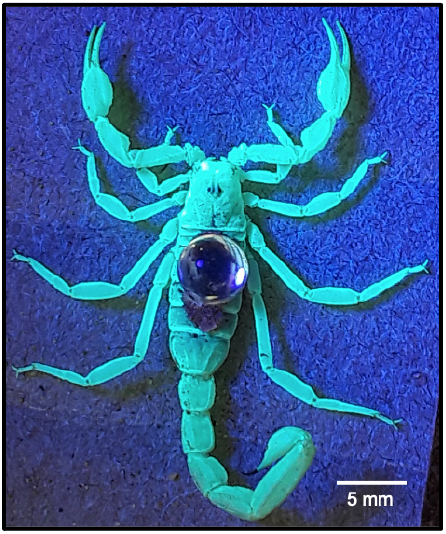
Animal with attached crystal. A *P. utahensis* female is photographed under UV light with a 5 mm round cab crystal affixed via double sided tape to its dorsal mesosoma.

Before each recording, we created a mound either in the center of an arena or offset from the center in various positions. The video monitoring system was then set to record for a given length of time. Finally, a crystal-equipped animal was introduced near the wall of its designated arena and the curtains were completely closed around the front of the set-ups using binder clips.

### Inducing learning walks without mounds

We also induced scorpions to occupy burrows in the center of the arenas without pre-made mounds. To do this, we added a thicker layer of sand (3000 ml) to the arena and placed a plastic ring (30 cm diam x 12.5 cm tall) in the middle. In the center of the plastic ring we partially buried a small paper slip (formed by removing the base of a Dixie 3 ounce bath cup). We misted over the top of the slip with 5 squirts (∼4 ml) of water to provide additional structural support to induce the scorpion to dig within this smaller arena. We then placed a scorpion in the center ring in the late afternoon and used video recording and MATLAB to track the animal’s movements for 18-22 hours. The plastic ring was removed the following afternoon if the scorpion was found inside or near the burrow. If the scorpion failed to accept the burrow, the smaller ring was left in place, the burrow region was misted with three additional squirts of water, and the animal was given an additional night to burrow. After the plastic ring was removed, we continued recordings to track the animal’s movements throughout the large arena for an another night.

### Analysis

We wrote various MATLAB scripts to analyze our behavioral data. We used a frame-by-frame subtraction method coupled with centroid plotting to automatically track the X-Y coordinates of the scorpion locations during our videos. We then used the Pythagorean theorem to calculate the distance walked and used the video frame capture rate to determine velocity. We also made time-lapsed videos that plotted the current animal position along with the three previous positions to create a stardust effect, which efficiently revealed instances of the animal’s initial burrowing. Once the initial digging was identified, we then hand plotted (for increased accuracy) the animal’s subsequent movements until we were confident that the animal had resumed its exploratory behavior or remained in the burrow for a prolonged period.

## RESULTS

### Activity plots

In all, we tracked 23 different animals, some for multiple evenings, for a total of nearly 1500 hours of video. During our trials, the animals spent most of their time walking along the walls of the arena but also made many forays across the arena’s interior. All night plots of animal movements (Fig 4) show a lot of activity, including concentrated movements around the central mound.

**Fig. 4.**
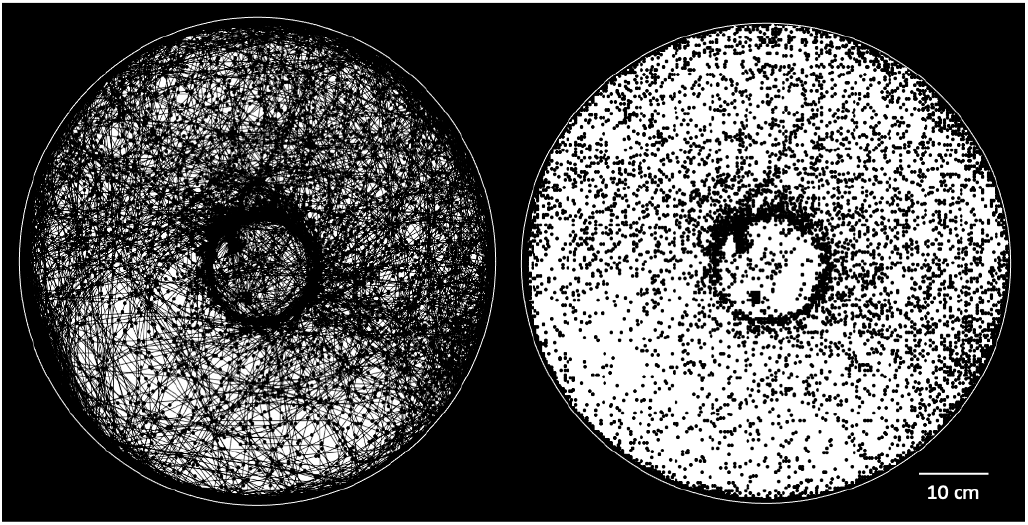
Long term activity plot. Shown is an example of an all-night video with the animal’s position plotted every second via a MATLAB script. The animal’s paths are shown by connecting the points with line segments in the left plot; the segments are excluded in the right plot.

### Burrow formation

As in our pilot studies (see supplemental materials), the animals in these trials readily dug burrows in the central mounds. Most of the initial digging occurred toward the end of the dark period or soon after the lights turned on. A sampling of some of the burrows we observed are shown in Fig 5, along with an example of a natural scorpion burrow filmed on the wildlife refuge. A short video clip of a scorpion digging its burrow is provided in the supplemental material (video S1).

**Fig. 5.**
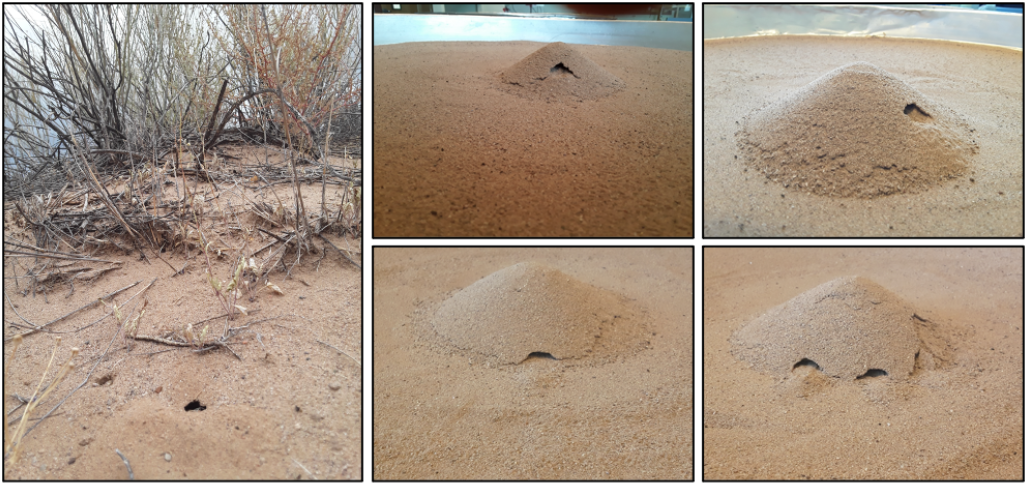
Scorpion burrows in nature and lab. The photo at left is an example of a scorpion burrow next to one of the field station’s trails. The four photos at right are examples of burrows we saw in our trials.

### Activity patterns

Figure 6 shows activity plots by hour for an animal we monitored for seven consecutive days. Over the seven days the animal walked 4,415 m for an average of 631 m per night. Tracking this animal’s average distance walked by hour of the day showed a consistent pattern of behavior (Fig 6, top histogram) with the highest activity soon after the arena lights turned off in the evening and soon before or just after the lights turned on in the morning. This pattern was also evident when the activities of all animals were pooled (Fig 6, bottom histogram).

**Fig. 6.**
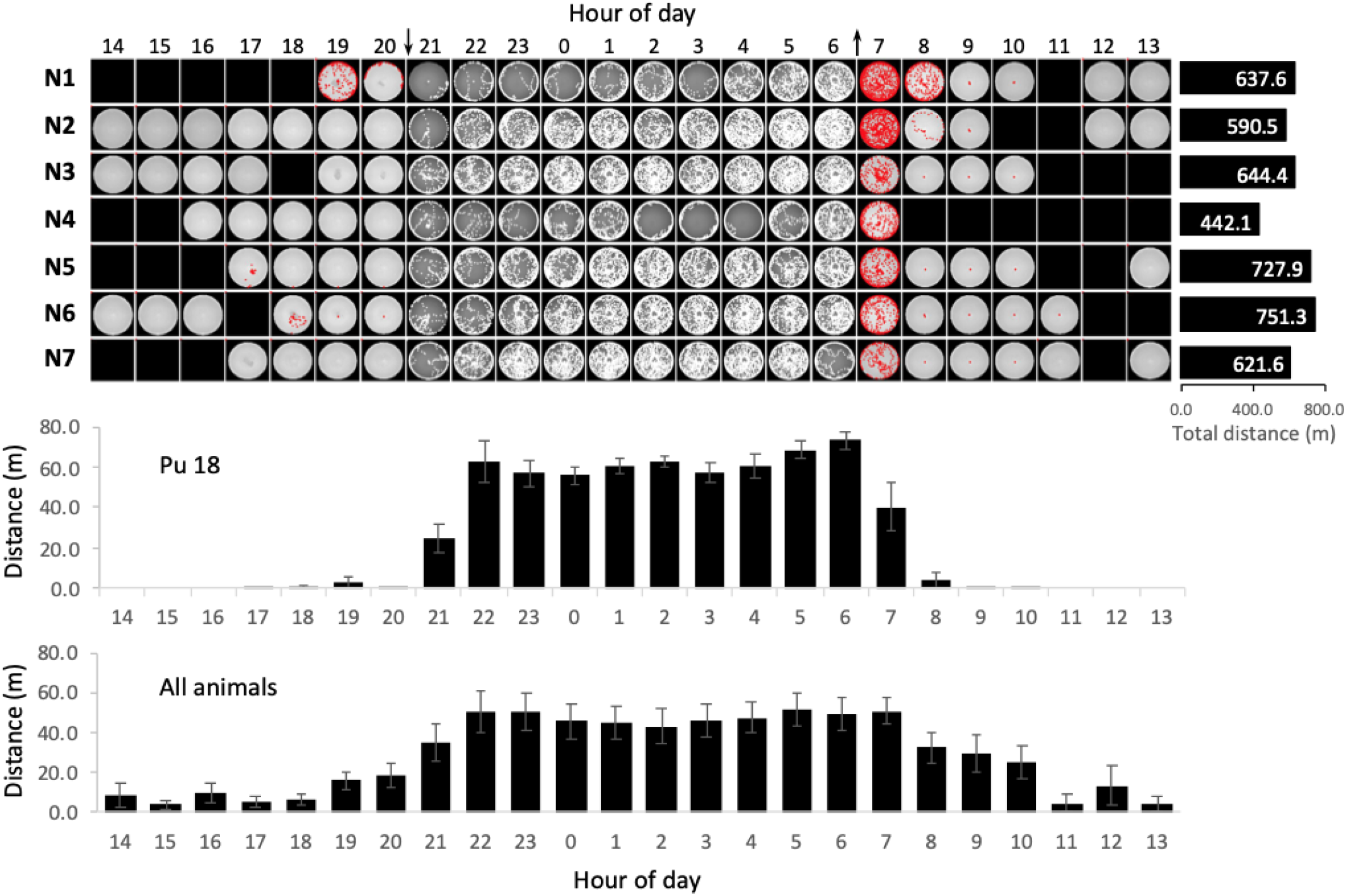
Animal activity patterns. At top are plots for a single animal over seven consecutive days. The arrows at top indicate when the lights turned off (down arrow) and on (up arrow). The blank squares reflect periods when the video recording was paused. The distance walked each night (in meters) is shown in bars at right. The middle histogram (Pu 18) shows the mean distance walked by hour (±SD) for the animal depicted in the top plots. The bottom histogram is a summary of all the animals we tested (23 animals, ∼1500 hours of video sampled every 2 s).

### Signs of learning walks

An example of a typical learning walk following initial burrowing is shown in Fig 7; a video of this walk is provided in the supplemental materials (video S2). Figure 7 also shows how we processed video showing the looping excursions. We hand-marked the position of the burrow and used the Pythagorean theorem to plot the distance of the animal from the burrow over time (top graph). We also plotted the animal’s instantaneous velocity by time during the walk (second graph). Next we plotted the distance from the burrow against the cumulative path length (third graph) and marked each return to the burrow. Finally we superimposed each of these individual loops by plotting the start of each at the origin (bottom graph).

**Fig. 7.**
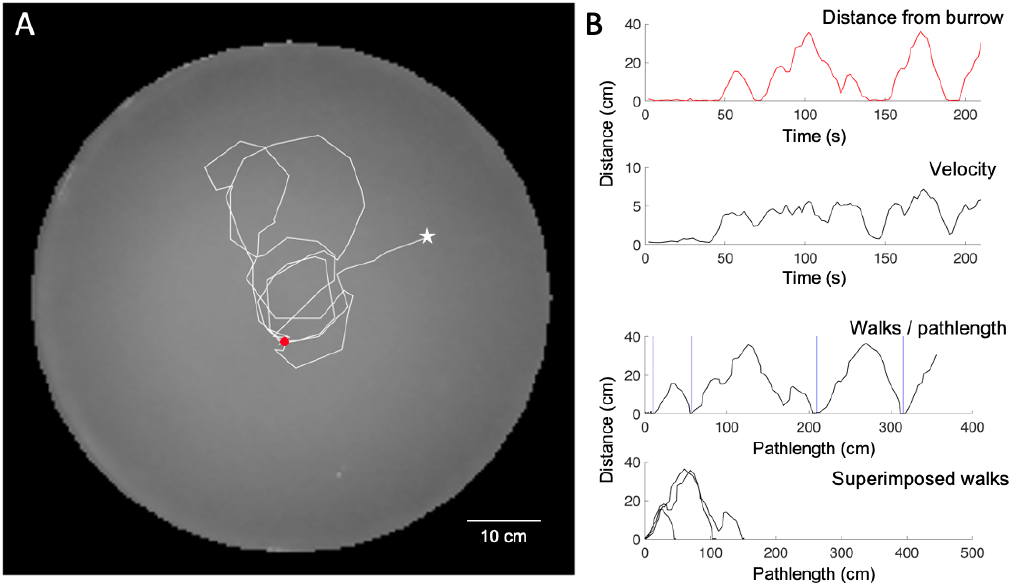
Sample learning walks and processing. (A) Approximately 200 s of an animal’s initial learning walk is plotted (red dot = burrow; white star = end of walk). (B) We processed the walks by first plotting distance from the burrow by time (upper graph) and the velocity of the animal by time (second graph). We then transformed the data to distance from the burrow over path length (third graph) and sliced out the walks based on each return to the burrow (vertical blue lines) and superimposed the walks in the bottom graph.

Of the 23 animals we monitored, 18 showed looping walking behavior immediately after the first signs of burrow digging. For the other 5 animals, the video resolution either did not allow accurate detection of digging behavior, or burrow formation happened outside the period of video monitoring. Fig 8A shows all of the initial learning walks that we encountered for these 18 animals along with the processing described in Fig 7. In all, 80 looping excursions away and back to the burrow were identified for all of the animals and these are superimposed in the graph of Fig 8B. The number of loops observed per animal varied from 1 to 10 and averaged 4.4 ±2.5 (SE). The average duration of the initial learning walks was 348.9 s ±47.9 (SE) and the average distance covered was 505.6 cm ±74.6 (SE). We determined the average velocity of each animal’s initial learning walk by dividing the distance covered by the duration of the walk. The average velocity of these walks was 1.7 cm/s ±1.4 (SE).

**Fig. 8.**
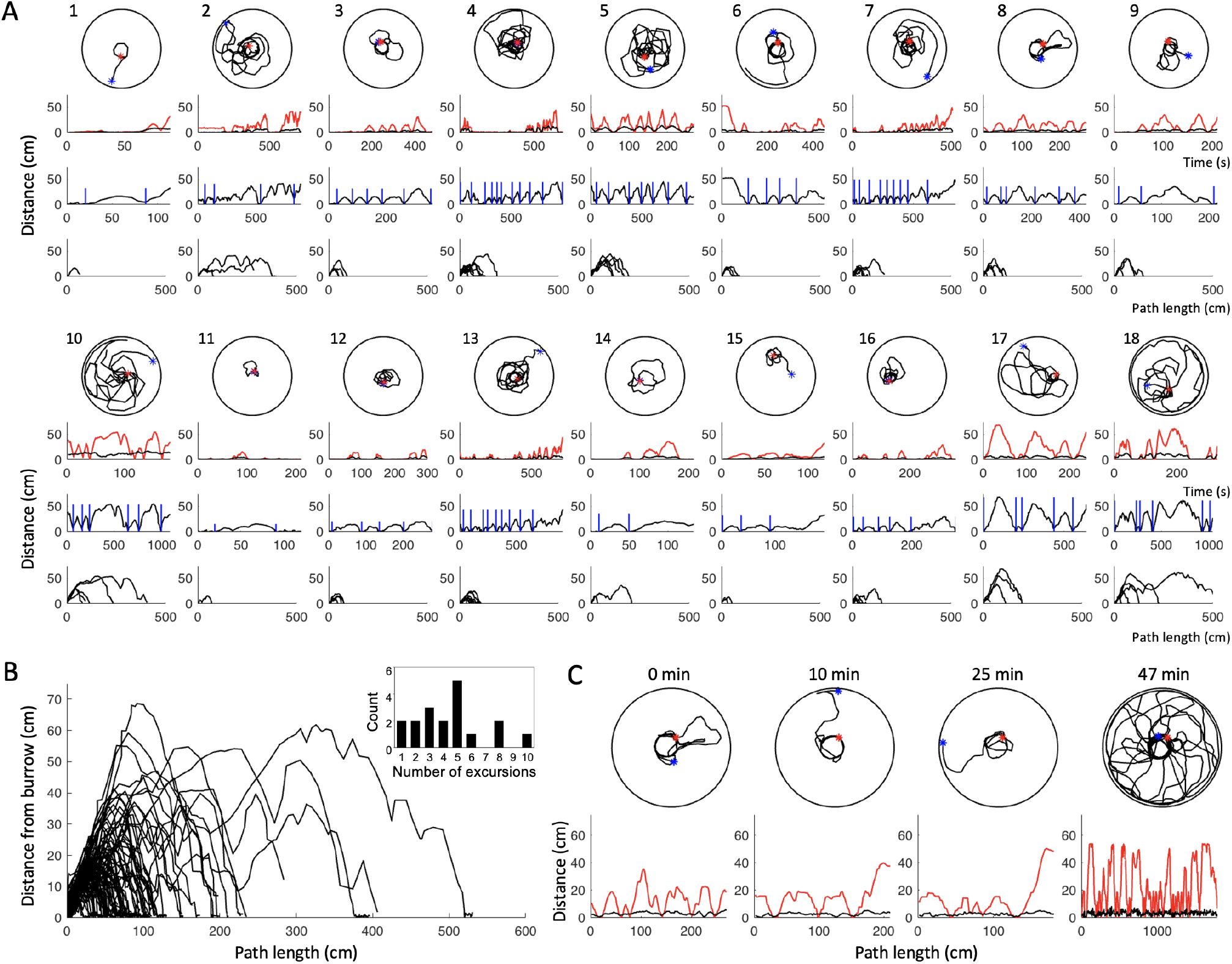
Learning walks. (A) All initial learning walks observed in the study (18) are plotted and processed as described in Fig 7. (B) All walks identified in the trials (total = 80) are superimposed. The inset shows the number of looping excursions observed during the initial walks. The number of loops ranged from one to ten and averaged 4.4 per animal. (C) The animals often made subsequent looping excursions later in the recordings. This example shows additional walks 10, 25, and 47 minutes after the first set of walks.

The focus of this study was on capturing the first occurrence of putative learning walk behavior immediately after the initial signs of burrow digging. However, the animals displayed many subsequent looping routes later in the videos. Some of these became elaborate and encompassed all parts of the arena. One such example is shown in Fig 8C where bouts of looping excursions occurred 10, 25, and 47 minutes after the initial set.

### Learning walks without a mound

We tried to reduce the possible visual or tactile influence of the sand mound by inducing several animals (n=9) to adopt burrows in level sand in the middle of the arena. Figure 9A shows three examples the first set of looping excursions that animals made after their first return to the burrow in the middle of the arena. Figure 9B shows an example of subsequent bouts that occurred later in the recording.

**Fig. 9.**
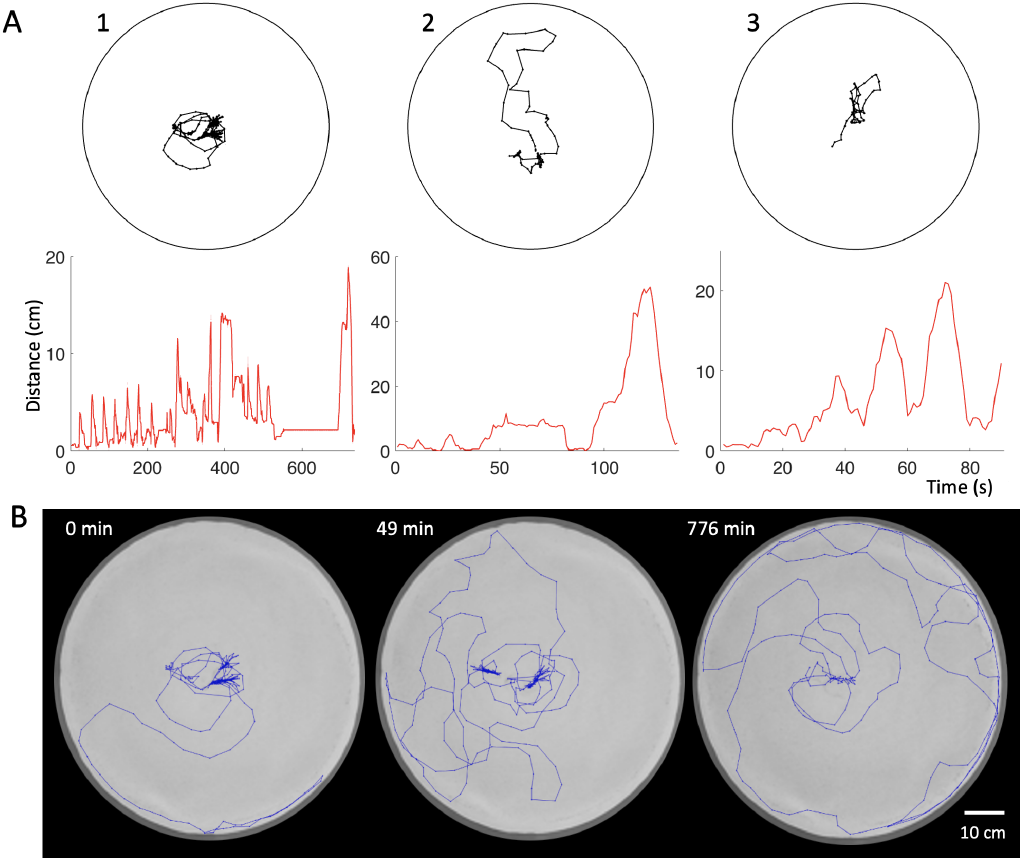
Learning walks without a mound. (A) Examples of the initial set of looping excursions for three animals that were induced to burrow in level sand in the middle of the arena. (B) Example of multiple bouts of looping excursions for animal 1 in A. At left is the initial set (time 0); the other two sets were detected at 49 and 776 min after the initial bout. The bouts were 21, 19, and 11 min long, respectively.

## DISCUSSION

Our findings are clear. Essentially all animals that made their own burrows in the middle of our laboratory arenas executed looping walks immediately after their first signs of digging. We found similar looping excursions whether we induced the animals to burrow in a small sand mound or in level sand in the middle of the arenas. This is the first report of learning walks in scorpions.

Learning walks are consistent with the navigation by chemotextural familiarity hypothesis. In this view, the putative learning walks could be an innate behavior that allows scorpions to acquire home-directed tastes and touches of the substrate near their burrow for subsequent retracing (Gaffin and Brayfield, 2017; Musaelian and Gaffin, 2020). This idea is similar to that proposed for familiarity navigation in desert ants (Baddeley et al., 2011; 2012; Gaffin and Brayfield, 2016), but instead of acquiring panoramic visual glimpses via compound eyes, the scorpion pectines act as a local sensor that acquires matrices of chemo-textural information from the substrate beneath the animal. This local sensor approach was used in a computer simulation that used straight down views of Earth satellite images to navigate (Gaffin et al., 2015) and has been applied to simulations of scorpion navigation (Gaffin and Brayfield, 2017; Musaelian and Gaffin, 2020).

In hymenopterans, learning walks and learning flights appear to help the animals learn home-related features of the landscape (Degen et al., 2016; Collett and Zeil, 2018). These walks or flights, if directed in various directions from the hive or nest, also keep an animal from overshooting its home when following a longer home-bound vector (such as generated through path integration). This is because the scenes, tastes, and touches beyond the nest are unfamiliar unless there was a way to acquire a repertoire of home-directed scenes which bend back to the starting point. Indeed, the addition of artificial learning walks to a computer simulation improved the homing accuracy of artificial agents navigating by scene familiarity (Baddeley et al., 2012).

Other observations from this experiment also appear in line with the chemo-textural familiarity idea. First, we tried rotating a few of the arenas midway through a second evening, when the animal was away from its burrow (the curtain and other visual features of the setup remained in place). An example is shown in Fig 10A. The animal repeatedly returned to the rotated burrow (instead of the position prior to rotation). This is not surprising since the burrow position did not change relative to the animal’s position. Future experiments should try lifting the animal before rotating the arena, replacing the animal in a new position relative to the burrow. We would expect the animal would still use learned substrate information to return to the position of the rotated burrow.

**Fig. 10.**
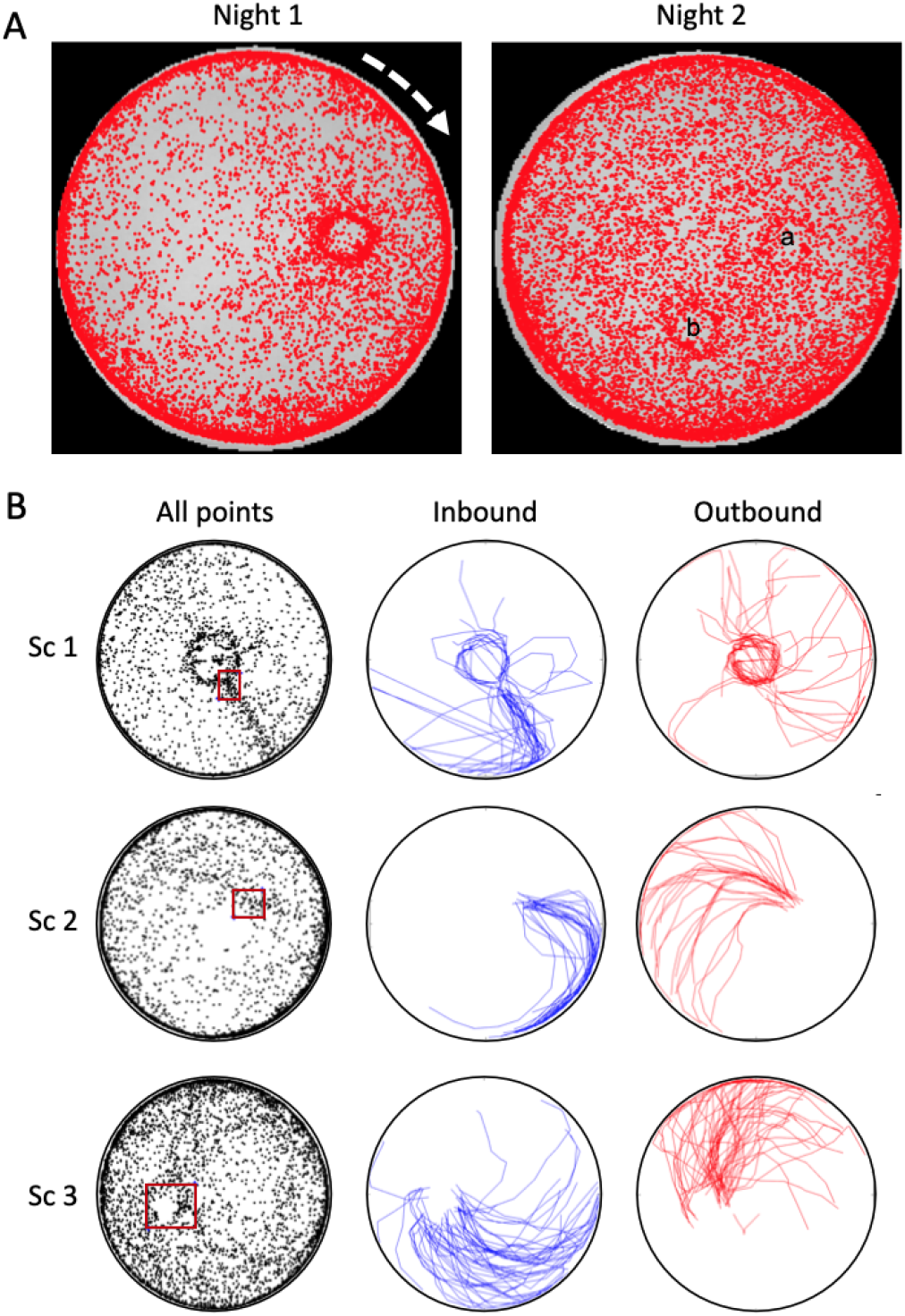
Additional behavioral observations. (A) Animal behavior after arena rotation. Two nights of activity are shown. In night 1, the arena was main-tained in its original orientation. Midway through night 2, the arena was rotated clockwise 90°. The position of the burrow can be detected in the night 2 plot both before (position “a”) and after (position “b”) the rotation. (B) Shown are some all-night plots where we drew a small rectangle around the burrow region and used MATLAB to plot the previous 20 seconds and sub-sequent 20 seconds of movement to that area. The animals used consistent inbound paths, which were different from the outbound paths.

We also saw some interesting patterns when we analyzed inbound and outbound paths relative to burrow location (Fig 10B). We digitally placed a rectangle around the position of the animal’s burrow after movement coordinates had been gathered for an entire evening. We then used MATLAB to plot the twenty seconds of movement prior to the animal entering the rectangle (“Inbound” paths) and the twenty seconds of movement after the animal exited the rectangle (“Outbound” paths). Interestingly, the animals tended to follow consistent and concentrated inbound paths that were strikingly different from their more dispersed outbound paths. These movement patterns suggest that previously learned features may guide animals along consistent homedirected routes.

In the Introduction, we noted that adequate sensor and environmental complexity is necessary for animals or agents navigating via familiarity to avoid being confused by similar scenes, tastes, or textures in multiple locations. This trade-off has been examined in various vision-based simulations (Gaffin et al., 2015; Gaffin and Brayfield, 2016) and in a navigation simulation modeled on scorpion pectines (Musaelian and Gaffin, 2020).

Our estimates of the pattern detection capacity of scorpion pectines are informed by electrophysiological studies showing that peg sensilla responded similarly to a variety of chemicals presented to the pore at the peg tip (Knowlton and Gaffin, 2009; 2010; 2011a;b; Gaffin and Walvoord, 2004). Based on these data it has been estimated that the pectines can conservatively detect from 10^12^ to 10^40^ different patterns (Gaffin and Brayfield, 2017). Further, neurons in peg sensilla interact synaptically (Gaffin and Brownell, 1997a; Foelix and Müller-Vorholt, 1983; Melville, 2000; Gaffin, 2002) which appears to reduce sensory adaptation through a local feedback loop and may improve information fidelity for navigation (Gaffin and Shakir, 2021).

Quantifying the chemo-textural complexity of the scorpion’s sand substrate however, is difficult. Proxies of the textural information available on the surface of a fine sand substrate (and at dimensions germane to the packing densities of the peg sensilla matrices) have been generated by photographing multiple images of sand through a dissecting microscope while projecting light from the side to produce pronounced shadows (Gaffin and Brayfield, 2017; Musaelian and Gaffin, 2020). Knowing the nature of the chemical milieu that occurs naturally on sand grains is still more challenging. Studies of scorpion responses to pheromone deposits suggest the chemicals may stably adhere to the sand grains and remain viable for scorpion sensory detection for days to weeks (Gaffin and Brownell, 1992; Taylor et al., 2012). It seems safe to suggest that decaying organic matter, animal deposits, and numerous other processes leave hundreds of residual chemicals on the sand in varying concentrations, creating enormous chemical complexity. Simply put, the peg matrices and substrate appear suitably matched in complexity.

Many additional studies are needed to build on the results presented in this study. For example, while we ran our experiments under IR cameras and attempted to exclude as much extraneous light as possible, scorpion eyes are sensitive to starlight levels of light (Fleissner and Fleissner, 2001). As such, it is crucial to repeat these tests using animals whose eyes have been thoroughly blocked with blindfolds. The arena lights should also be smoothly dimmed and brightened to simulate natural sunset and sunrise conditions. Other experiments should consider disrupting the sand around the burrow after bouts of walks have occurred to see if looping behavior intensifies relative to baseline levels without disruption. In addition, disruption of the sand while the animal is away from its burrow would be useful for assessing the use of home-directed substrate information. Tests also need to systematically alter the rotation of the arena relative to the curtain and the laboratory to control for visual and geocentric cues. Finally, we think it would be interesting to look for signs of learning walks in other long range navigating arachnids, such as amblypygids, that have substantial chemoand mechanosensory attributes (Hebets, 2002; Hebets et al., 2014).

## Acknowledgements

We thank George Martin for assistance with our behavioral set-up, Jacob Sims and Joe Bradley for help collecting animals, the UNM Sevilleta Field Station and personnel for lodging and research support and the Sevilleta LTER, especially Kathy Granillo (refuge manager), for access to field sites. We also thank Alexis Merchant and Hannah Peeples for reviewing the manuscript. Finally, we thank Sandra Doan and Gail Goodson of the Laboratory Animal Resources facility for help establishing our animals and behavioral assays at the University of Oklahoma.

## Contribution

Conceptualization: D.G.; Methodology: D.G., M.H., M.M.; Validation: D.G.; Formal analysis: D.G., M.M.; Investigation: D.G., M.H., M.M.; Resources: D.G.; Data curation: D.G.; Writing original draft: D.G.; Writing review & editing: M.H., D.G., M.M.; Visualization: D.G.; Supervision: D.G.; Project administration: D.G.; Funding acquisition: D.G..

## Supplementary

Supplementary figures and videos can be found here.

